# Behavioural diversity of bonobo prey preference as a potential cultural trait

**DOI:** 10.1101/2020.06.02.130245

**Authors:** L. Samuni, F. Wegdell, M. Surbeck

## Abstract

The importance of cultural processes to behavioural diversity, especially in our closest living relatives, is central for revealing the evolutionary origins of human culture. Whereas potential cultural traits are extensively investigated in chimpanzees, our other closest living relative, the bonobo, is often overlooked as a candidate model. Further, a prominent critique to many examples of proposed animal cultures is premature exclusions of environmental confounds known to shape behavioural phenotypes. We addressed these gaps by investigating variation in prey preference expression between neighbouring bonobo groups that associate and share largely overlapping home ranges. We find specific group preference for duiker or anomalure hunting that are otherwise unexplained by variation in spatial usage of hunt locations, seasonality or sizes of hunting parties. Our findings demonstrate that group-specific behaviours emerge independently of the local ecology, indicating that hunting techniques in bonobos may be culturally transmitted. We suggest that the tolerant intergroup relations of bonobos offer an ideal context to explore drivers of behavioural phenotypes, the essential investigations for phylogenetic constructs of the evolutionary origins of culture.

## Introduction

Humans and other social animals exhibit a diversity of behavioural phenotypes attributed to genetic and/or social (i.e., culture) evolutionary processes influenced by the environment (Castro and Toro, 2004; Van Schaik et al., 2003; Whiten, 2017). While culture is identified as a pivotal selective process in human evolution (Boyd and Richerson, 1995; Castro and Toro, 2004; Creanza et al., 2017), its relative contribution to shaping the behavioural diversity observed in non-human animals, including our closest living relatives, remains debated. For instance, in comparison to the other great ape species, little is known about potential cultural traits in bonobos (*Pan paniscus*) (Whiten, 2017), thereby limiting phylogenetic comparisons.

Culture is defined as group-specific behavioural patterns acquired through some form of social learning (Laland and Janik, 2006). There is ample evidence that some foraging techniques are socially learned (e.g., primates (Whiten and van de Waal, 2018); cetaceans (Mann et al., 2012); carnivores (Thornton and Raihani, 2008)) and therefore represent good candidates for cultural traits. However, to distinguish whether social processes contribute to the emergence of behavioural phenotypes, it is essential to quantify ecological variation and account for its influence on behaviour expression, a challenging endeavour in wild settings.

Our closest living relatives, bonobos and chimpanzees, hunt a variety of species across groups and populations (Gilby et al., 2015; Hobaiter et al., 2017; Hohmann and Fruth, 2008; Sakamaki et al., 2016; Samuni et al., 2018; Wakefield et al., 2019). However, it remains unclear whether this diversity is independent of large or even small-scale ecological variation in the distribution of prey species (Hobaiter et al., 2017; Sakamaki et al., 2016). Accounting for potential local ecological drivers is methodologically challenging in chimpanzees, a territorial species where each group predominantly occupies unique non-overlapping areas (Mitani et al., 2010; Samuni et al., 2017). In contrast, the tolerant intergroup relations of bonobos (Furuichi, 2020; Lucchesi et al., 2020) permit a context in which different behaviours are expressed by individuals of different groups in the same place and at the same time. Here, we investigate variation in bonobo predation patterns of two groups (Ekalakala and Kokoalongo) at the Kokolopori Bonobo Reserve. The groups share an extensive home range overlap (65% kernel and 80% maximum convex polygon overlaps; Fig. 1) and regular gene flow, thereby reducing ecological and genetic influences as an explanatory variable for inter-group differences in behavioural expressions. Specifically, we tested whether variation in prey preference between the two bonobo groups is explained by a) environmental variables, such as area usage and seasonality, and/or b) social factors, such as the number of hunters and group identity.

**Figure 1.**
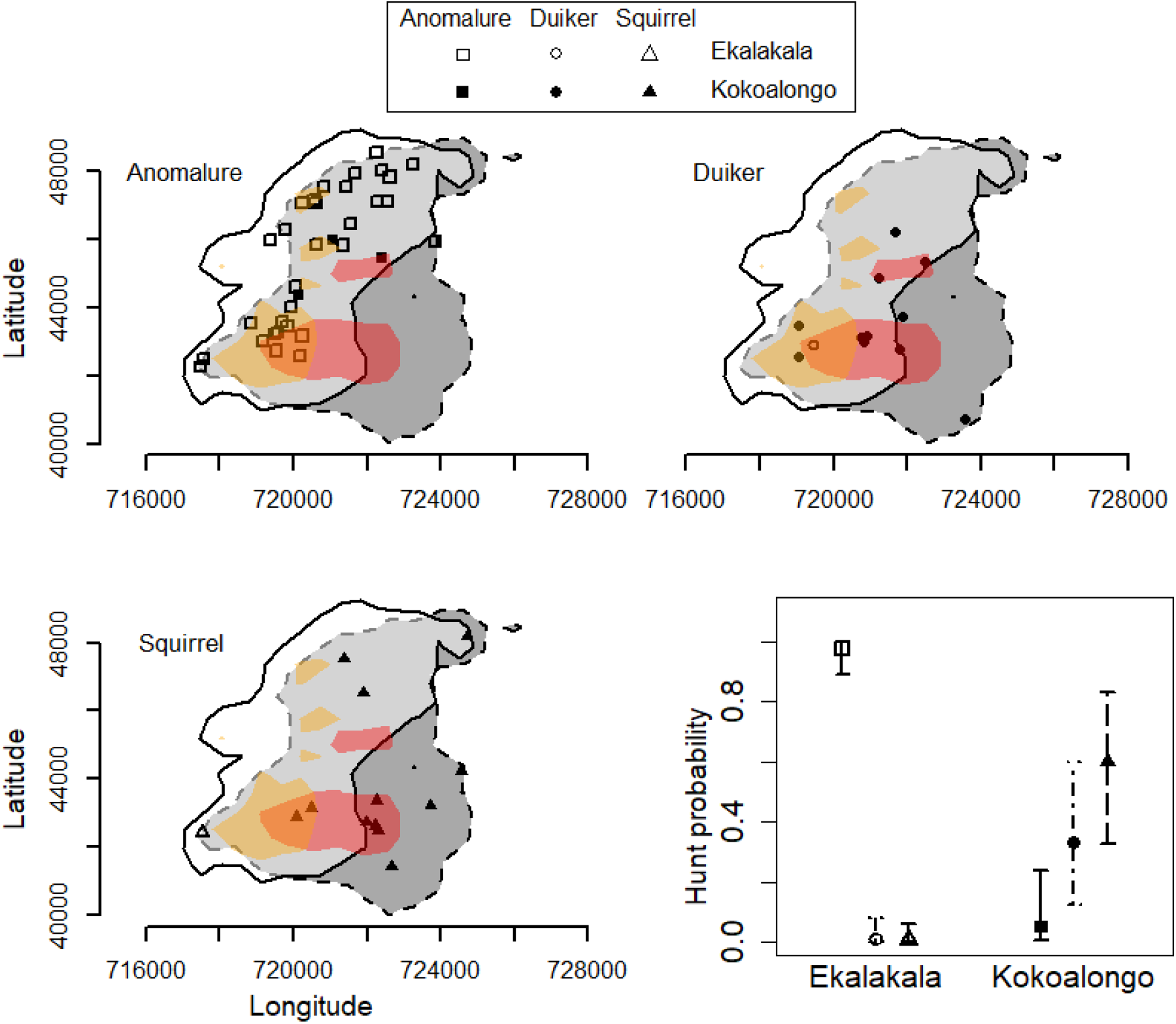
Predation patterns in Kokolopori bonobos. Hunting locations of the three prey types: anomalure (square; top left), duiker (circle; top right), and squirrel (triangle; bottom left) in relation to the 95% Kernel usage area of Ekalakala (white polygon with solid border) and Kokoalongo (dark grey polygon with dashed border) and 50% Kernel usage area (Ekalakala in yellow, Kokoalongo in red). The overlapping 95% kernel area between Ekalakala and Kokoalongo is depicted in light grey. Also depicted are hunt probabilities of the different prey types between Ekalakala and Kokoalongo (bottom right) as obtained from the BR model.

## Methods

### (a) Study site and data collection

We investigated behavioural diversity between two fully habituated bonobo groups (Ekalakala and Kokoalongo, habituated since 2007) at the Kokolopori Bonobo Reserve, Democratic Republic of Congo (N 0.41716°, E 22.97552°; (Surbeck et al., 2017)). We conducted full day party follows of the bonobo groups (1102 and 931 observation days in Ekalakala and Kokoalongo, respectively) and documented all occurrence hunting behaviour (here defined as capture of mammalian prey). All prey types were captured across most months, and both during the dry (June-August and December-February) and wet (March-May and September-November) seasons. Hunt participants were almost exclusively adult (>10 yrs) individuals, and both sexes were observed to participate. Adult group sizes fluctuated during the study between 9-11 adult individuals in Ekalakala and 16-24 adult individuals in Kokoalongo due to several deaths and migration events.

### (b) Home range utilization distribution

We recorded data on party locations at one-minute intervals using a GPS (Garmin© 62). We constructed home range utilization distributions of the bonobo groups using kernel density estimates (Worton, 1989). The home range (95% kernel) of the two groups between August 2016 and December 2019 was: Ekalakala – 35km^2^, Kokoalongo – 40km^2^, and the overlapping area encompassed 64% and 66% of the home rages of Ekalakala and Kokoalongo, respectively.

Habitat structure and spatial distribution of prey species has been used as explanations for variation in hunting behaviours (Hobaiter et al., 2017; Sakamaki et al., 2016). However, as our data originate from two groups with extensive home range overlap, the explanatory power of these drivers is minimized. Nonetheless, we can evaluate intra-range variation in local ecology by accounting for relative home range usage across the groups. To do so, we assigned each hunt with two kernel usage values, one constituting the kernel usage of the group that hunted (*hunt group*) and the other constituting the kernel usage of the group that did not hunt (*other group*). We used the values to calculate a score of ‘usage difference’ (i.e., *other group* - *hunt group*; ranging between −50 and 86; mean ± sd: 20.19 ± 26.10). Higher scores reflected an area that is more predominantly used by the group that hunted.

### (c) Statistical analysis

We applied a Bayesian Regression model with prey type as a categorical response and logit link function to examine the influence of environmental (area usage and seasonality) and social (group identity and presence of potential hunters) factors on prey preference expression. We fitted the model in R (version 3.6.1 (R Core Team, 2016)) using the function *brm* of the R package ‘brms’ (Bürkner, 2017) and the default flat priors. As predictors, we included the following environmental factors: a) ‘usage difference’ score as described above, and b) a seasonal temporal term, by including the sine and cosine of the Julian dates of the hunts converted into a circular variable (Stolwijk et al., 1999). Additionally, we included the following social factors: a) group identity and b) adult party size recorded at 30-minute intervals. We did not differentiate between the number of adult males versus adult female party members as we do not expect it to have a difference on prey acquisition (both were regular hunt participants in our study).

We ran 2,000 iterations over four MCMC chains, with a ‘warm-up’ period of 1,000 iterations per chain leading to 4,000 usable posterior samples (Bürkner, 2017). Visual inspection of all MCMC results revealed satisfactory Rhat values (< 1.01; (Gelman et al., 2013)), no divergent transitions after warmup, and stationarity and convergence to a common target, suggesting that our results are stable. We report the estimate (mean of the posterior distribution) and the 95% credible intervals (CI_95%_) indicating the strength of the effects. For estimate comparability and to ease model convergence, we z-transformed ‘usage difference’ and party size before fitting the models. The data reported in this paper are available as a supplementary dataset.

## Results

Between August 2016 and January 2020 we observed 59 successful captures and consumption of mammals by the bonobos, including anomalure, duiker, and squirrel species (Table 1). During a six months period (Jul-Dec 2019) we also documented 11 unsuccessful hunt attempts on duiker and anomalure in both groups (duiker-N_Ekalakala_ = 2, N_Kokoalongo_ = 2; anomalure-N_Ekalakala_ = 4, N_Kokoalongo_ = 3). Overall, we observed all Ekalakala and 84% of Kokoalongo adult group members (100% if considering only individuals that were present for the entire study period) participating in hunts.

**Table 1.** Successful hunts in Ekalakala and Kokoalongo between August 2016-Jan 2020

A. occurrence of successful hunts
| Group | Duiker species<br>(*Philantomba monticola*,<br>*Cephalophus castaneus*) | Squirrel species<br>(*Funisciurus congicus*) | Anomalure species<br>(*Anomalurus derbianus*,<br>*Anomalurus beecrofti*) |
| --- | --- | --- | --- |
| Ekalakala | 1 | 1 | 31 |
| Kokoalongo | 11 | 12 | 3 |

Group home ranges extensively overlapped and most anomalure and duiker hunts occurred within the overlapping areas (94% of anomalure and 83% of duiker hunts), as opposed to 46% of squirrel hunts (Fig. 1). The groups engaged in frequent and prolonged intergroup associations (31% of observation days), and nine of the hunts (5 duiker, 3 anomalure, 1 squirrel) occurred during intergroup encounters and involved between-group meat sharing. Due to the cohesiveness of bonobo groups (Hohmann and Fruth, 2002), the conspicuous nature of anomalure and duiker hunting (e.g., distress calls of duikers), and since the acquisition of meat often attracts individuals to hunting areas (Samuni et al., 2018), we are confident that we observed most anomalure and duiker feeding events. However, as the hunting and feeding of squirrel is often quiet and solitary, we may have underestimated this type of hunting.

Kokoalongo bonobos were more likely to capture duiker (estimate = 7.09, CI_95%_ = [3.71, 12.00]; Fig. 1, Table 2) and squirrel species (estimate = 7.49, CI_95%_ = [4.14, 12.17]), and were less likely to capture anomalure species in comparison with Ekalakala. The same pattern persisted during intergroup encounters (once we observed anomalure captured by a Kokoalongo female after a hunt by Ekalakala individuals). We found that group-specific prey preferences were independent from potential local spatial and temporal ecological variation. Overall, more than 80% of all hunts occurred in overlapping areas (95% kernel), and neither utilization differences of specific hunt locations (reflecting varying opportunities to encounter prey species) nor potential annual seasonal variation strongly affected phenotypic variation in prey types captured (Table 2). Variation in prey preference can also arise from between-group difference in sizes of association parties, or presence of certain specialized hunters. However, the number of adult individuals present during hunts (i.e., available hunters; mean ± sd: Ekalakala - 7.9 ± 1.6; Kokoalongo - 8 ± 4.3; encounter - 14.5 ± 7.4) had no strong effect on prey outcome (Table 2). Further, we observed 16 different individuals (four males and 12 females) catching prey, encompassing 72% of Ekalakala and 40% of Kokoalongo group members. These percentages are likely an underestimation of the overall number of individuals who captured the prey, as their identity was recorded for less than half of all hunts. Finally, our results are likely independent from genetic variation, as low genetic differentiation is expected (Schubert et al., 2011) mainly due to regular gene flow attributed to female migration between Ekalakala and Kokoalongo.

**Table 2.** Bayesian Regression model results of the effect of group identity, number of available hunters and ecological variation on prey species captured (^1^anomalure and ^2^Ekalakala as reference categories).

| Coded level |  | Estimate | SE | 95% CI |
| --- | --- | --- | --- | --- |
| Duiker<sup>1</sup> | Intercept | -4.49 | 1.48 | -7.91, -2.26 |
|  | **Group (Kokoalongo<sup>2</sup>)** | **7.09** | **2.13** | **3.71, 12.00** |
|  | Available hunters | 0.72 | 0.57 | -0.34, 1.95 |
|  | Usage difference | 0.59 | 0.65 | -0.59, 1.96 |
|  | Sine of Date | 1.33 | 1.06 | -0.55, 3.59 |
|  | Cosine of Date | 0.45 | 1.15 | -1.69, 2.82 |
| Squirrel<sup>1</sup> | Intercept | -4.58 | 1.39 | -7.76, -2.45 |
|  | **Group (Kokoalongo<sup>2</sup>)** | **7.49** | **2.07** | **4.14, 12.17** |
|  | Available hunters | -0.07 | 0.63 | -1.27, 1.21 |
|  | Usage difference | 0.86 | 0.65 | -0.31, 2.24 |
|  | Sine of Date | 1.26 | 1.07 | -0.64, 3.68 |
|  | Cosine of Date | 1.00 | 1.14 | -1.11, 3.37 |

## Discussion

Overall, our findings demonstrate that social processes rather than local ecological variation predicted group-specific prey acquisition in Kokolopori bonobos, providing strong indication of cultural traits in wild bonobos. Observed differences in prey species preferences may arise if different techniques are required to locate and capture them. Duiker and squirrel hunting are either strictly terrestrial (duiker) or arboreal (squirrel) activities, which appear opportunistic and commonly involved a single individual hunter (more so for squirrel hunting). Conversely, anomalure hunting required the engagement of several group members, during which the bonobos employed both terrestrial and arboreal positions. At this stage it is unclear which forms of learning such hunting techniques in bonobos may necessitate (e.g., emulation, stimulus enhancement, or imitation). However, as successful prey acquisition can be influenced by social learning (Mann et al., 2012; Whiten and van de Waal, 2018), hunting techniques may emerge as a form of cultural transmission.

Another potential explanation for prey preference variation is the presence of individuals who act as social hunt catalysts (i.e., “impact hunters” (Gilby et al., 2015)). The ‘impact hunter hypothesis’ explains how joint hunt participation may emerge when hunts by single individuals are costly (e.g., risk of injury, low success probability). The hypothesis proposes that increased motivation of certain individuals to hunt (and potentially suffer initial costs) encourages others to join. For example, if certain group-members motivate social hunts (in this case anomalure), then the absence of such individuals in a group may lead to a bias towards solo hunting (in this case duiker and squirrel). Although apparent group-specific hunting behaviour may emerge as an artefact of the presence of impact hunters, patterns in our data indicate that this is unlikely to be the root of observed group differences. First, the absence of individualistic hunting (as apparent in Ekalakala) cannot be explained by the impact hunter hypothesis, as this hypothesis addresses social hunt occurrence. Second, although preliminary, similar frequencies of unsuccessful hunt attempt of anomalure in both groups, indicate that prey preference expression is likely driven by the group hunting technique efficiency rather than catalyst hunters. Additionally, as many adult group members participated in hunts and captured prey, collectively, our results indicate that we indeed document group, instead of individual, tendencies.

It is puzzling how such group differences would evolve and persist even when prolonged associations between Ekalakala and Kokoalongo should potentially promote intergroup social learning opportunities. Tolerance, at a degree that facilitates social learning in its various forms, is fundamental in converting innovations into transmitted traditions (Whiten and van de Waal, 2018). To improve “learning” gains, social learners should be selective in the timing of observations and their choice of “models” from whom to observe (Boyd and Richerson, 1995). Although the two groups associate for extended periods their intergroup relations are complex and unpredictable, characterized by a mixture of affiliative and agonistic exchanges, frequent fission-fusions and heightened arousal (Cheng et al., under review). Unpredictability of intergroup interactions is thus expected to hamper intergroup learning opportunities of certain skills which may require extensive time and effort to acquire (e.g., hunting techniques). Following group psychology predictions of ingroup bias and favouritism (Brewer, 1993), outgroup members may as well be less appealing “models” for learning. Together, inconsistent intergroup relations and in-group bias may explain how group-specific prey preferences persist despite numerous intergroup learning opportunities, indicating group bias and conformity. A by-product of divergent hunting techniques is reduced intergroup competition, which is likely adaptive, especially when groups share ranging zones. Thus, group-specific prey preferences in bonobos may have evolved as a form of microlevel niche differentiation that alleviates feeding competition.

Investigating the potential impact of culture on behavioural diversity in non-human animals is challenging due to the difficulties of estimating and accounting for local ecological variation as a driver of behavioural diversity. Challenges may even arise when behavioural variation appears between groups that occupy nearby but non-overlapping ranging areas. Bonobo social groups’ regular overlap in ranging area and tolerant interactions, offer fertile ground in which to explore whether variation in behavioural expressions occurs independently of spatial and temporal use of specific habitat locations. Here, by accounting and largely excluding potential local ecological variation, we provide strong indication for culturally transmitted subsistence hunting techniques in bonobos, informing on the evolution of behavioural diversity.

## Acknowledgements

We are grateful to the Bonobo Conservation Initiative and Vie Sauvage, especially Sally Coxe and Albert Lotana Lokasola, for their continuous support of this work. We thank the Ministry of Research of the Democratic Republic of the Congo for permitting the study, and the people of the villages of Bolamba, Yete, Yomboli and Yasalakose for granting access to their forest. We thank all the research assistants and local field assistants for their dedication and support in the field and for Erin Wessling and Catherine Hobaiter for their comments on this manuscript. This work is funded by the Max Planck Society and Harvard University.

The authors declare no competing interests, and all contributed to the methodology and writing of this work.

## Ethics Statement

The research presented here was non-invasive and approved by the Max Planck Society and the Ministry of Research of the Democratic Republic of the Congo. The authors declare that they have no conflict of interest.

## References

Boyd R, Richerson PJ. 1995. Why does culture increase human adaptability? Ethology and Sociobiology 16:125–143. doi:10.1016/0162-3095(94)00073-G

Brewer MB. 1993. Social Identity, Distinctiveness, and In-Group Homogeneity. Social Cognition; New York 11:150–164. doi:http://dx.doi.org.ezp-prod1.hul.harvard.edu/10.1521/soco.1993.11.1.150

Bürkner P-C. 2017. brms: An R Package for Bayesian Multilevel Models Using Stan. Journal of Statistical Software 80:1–28. doi:10.18637/jss.v080.i01

Castro L, Toro MA. 2004. The evolution of culture: From primate social learning to human culture. PNAS 101:10235–10240. doi:10.1073/pnas.0400156101

Cheng L, Lucchesi S, Mundry R, Samuni L, Deschner T, Surbeck M. under review. Variation in urinary cortisol levels indicates challenges of intergroup competition in wild bonobos.

Creanza N, Kolodny O, Feldman MW. 2017. Cultural evolutionary theory: How culture evolves and why it matters. PNAS 114:7782–7789. doi:10.1073/pnas.1620732114

Furuichi T. 2020. Variation in Intergroup Relationships Among Species and Among and Within Local Populations of African Apes. Int J Primatol. doi:10.1007/s10764-020-00134-x

Gelman A, Carlin JB, Stern HS, Dunson DB, Vehtari A, Rubin DB. 2013. Bayesian Data Analysis, 3 edition. ed. Boca Raton: Chapman and Hall/CRC.

Gilby IC, Machanda ZP, Mjungu DC, Rosen J, Muller MN, Pusey AE, Wrangham RW. 2015. ‘Impact hunters’ catalyse cooperative hunting in two wild chimpanzee communities. Phil Trans R Soc B 370:20150005. doi:10.1098/rstb.2015.0005

Hobaiter C, Samuni L, Mullins C, Akankwasa WJ, Zuberbühler K. 2017. Variation in hunting behaviour in neighbouring chimpanzee communities in the Budongo forest, Uganda. PLOS ONE 12:e0178065. doi:10.1371/journal.pone.0178065

Hohmann G, Fruth B. 2008. New Records on Prey Capture and Meat Eating by Bonobos at Lui Kotale, Salonga National Park, Democratic Republic of Congo. FPR 79:103–110. doi:10.1159/000110679

Hohmann G, Fruth B. 2002. Dynamics in social organization of bonobos (Pan paniscus) In: Boesch C, Hohmann G, Marchant L, editors. Behavioural Diversity in Chimpanzees and Bonobos. Cambridge University Press.

Laland KN, Janik VM. 2006. The animal cultures debate. Trends in Ecology & Evolution 21:542–547. doi:10.1016/j.tree.2006.06.005

Lucchesi S, Cheng L, Janmaat K, Mundry R, Pisor A, Surbeck M. 2020. Beyond the group: how food, mates, and group size influence intergroup encounters in wild bonobos. Behav Ecol 31:519–532. doi:10.1093/beheco/arz214

Mann J, Stanton MA, Patterson EM, Bienenstock EJ, Singh LO. 2012. Social networks reveal cultural behaviour in tool-using dolphins. Nature Communications 3:1–8. doi:10.1038/ncomms1983

Mitani JC, Watts DP, Amsler SJ. 2010. Lethal intergroup aggression leads to territorial expansion in wild chimpanzees. Current Biology 20:R507–R508. doi:10.1016/j.cub.2010.04.021

R Core Team. 2016. R: A language and environment for statistical computing. R Foundation for Statistical Computing, Vienna, Austria R Foundation for Statistical Computing, Vienna, Austria.

Sakamaki T, Maloueki U, Bakaa B, Bongoli L, Kasalevo P, Terada S, Furuichi T. 2016. Mammals consumed by bonobos (Pan paniscus): new data from the Iyondji forest, Tshuapa, Democratic Republic of the Congo. Primates 57:295–301. doi:10.1007/s10329-016-0529-z

Samuni L, Preis A, Deschner T, Crockford C, Wittig RM. 2018. Reward of labor coordination and hunting success in wild chimpanzees. Communications Biology 1:138. doi:10.1038/s42003-018-0142-3

Samuni L, Preis A, Mundry R, Deschner T, Crockford C, Wittig RM. 2017. Oxytocin reactivity during intergroup conflict in wild chimpanzees. PNAS 114:268–273. doi:10.1073/pnas.1616812114

Schubert G, Stoneking CJ, Arandjelovic M, Boesch C, Eckhardt N, Hohmann G, Langergraber K, Lukas D, Vigilant L. 2011. Male-Mediated Gene Flow in Patrilocal Primates. PLOS ONE 6:e21514. doi:10.1371/journal.pone.0021514

Stolwijk AM, Straatman H, Zielhuis GA. 1999. Studying seasonality by using sine and cosine functions in regression analysis. Journal of Epidemiology & Community Health 53:235–238. doi:10.1136/jech.53.4.235

Surbeck M, Coxe S, Lokasola AL. 2017. <NOTE>Lonoa: The Establishment of a Permanent Field Site for Behavioural Research on Bonobos in the Kokolopori Bonobo Reserve.

Thornton A, Raihani NJ. 2008. The evolution of teaching. Animal Behaviour 75:1823–1836. doi:10.1016/j.anbehav.2007.12.014

Van Schaik CP, Ancrenaz M, Borgen G, Galdikas B, Knott CD, Singleton I, Suzuki A, Utami SS, Merrill M. 2003. Orangutan Cultures and the Evolution of Material Culture. Science 299:102–105. doi:10.1126/science.1078004

Wakefield ML, Hickmott AJ, Brand CM, Takaoka IY, Meador LM, Waller MT, White FJ. 2019. New Observations of Meat Eating and Sharing in Wild Bonobos (Pan paniscus) at Iyema, Lomako Forest Reserve, Democratic Republic of the Congo. FPR 90:179–189. doi:10.1159/000496026

Whiten A. 2017. Culture extends the scope of evolutionary biology in the great apes. PNAS 114:7790–7797. doi:10.1073/pnas.1620733114

Whiten A, van de Waal E. 2018. The pervasive role of social learning in primate lifetime development. Behav Ecol Sociobiol 72:80. doi:10.1007/s00265-018-2489-3

Worton BJ. 1989. Kernel methods for estimating the utilization distribution in home-range studies. Ecology 70:164–168. doi:10.2307/1938423

